# ATG2A-mediated bridge-like lipid transport regulates lipid droplet accumulation

**DOI:** 10.1101/2023.08.14.553257

**Authors:** Justin L. Korfhage, Neng Wan, Helin Elhan, Lisa Kauffman, Mia Pineda, Devin M. Fuller, Abdou Rachid Thiam, Karin M. Reinisch, Thomas J. Melia

**Author notes:** Corresponding author, Thomas J. Melia.

## Abstract

ATG2 proteins facilitate bulk lipid transport between membranes. ATG2 is an essential autophagy protein, but ATG2 also localizes to lipid droplets (LDs), and genetic depletion of ATG2 increases LD numbers while impairing fatty acid transport from LDs to mitochondria. How ATG2 supports LD homeostasis and whether lipid transport regulates this homeostasis remains unknown. Here we demonstrate that ATG2 is preferentially recruited to phospholipid monolayers such as those surrounding LDs rather than to phospholipid bilayers. *In vitro*, ATG2 can drive phospholipid transport from artificial LDs with rates that correlate with the binding affinities, such that phospholipids are moved much more efficiently when one of the ATG2-interacting structures is an artificial LD. ATG2 is thought to exhibit ‘bridge-like” lipid transport, with lipids flowing across the protein between membranes. We mutated key amino acids within the bridge to form a transport-dead ATG2 mutant (TD-ATG2A) which we show specifically blocks bridge-like, but not shuttle-like, lipid transport *in vitro*. TD-ATG2A still localizes to LDs, but is unable to rescue LD accumulation in ATG2 knockout cells. Thus, ATG2 has a natural affinity for, and an enhanced activity upon LD surfaces and uses bridge-like lipid transport to support LD dynamics in cells.

## Introduction

ATG2 is a member of the VPS13 or repeating beta groove (RBG) family of lipid transport proteins (*1*), which are thought to function as hydrophobic bridges across which lipids can be transported at membrane contact sites (MCSs) (*2*). As an essential autophagy gene, ATG2 functions to support autophagosome membrane expansion (*3-8*). However, it has also been implicated in the regulation of lipid droplet (LD) homeostasis through numerous knockdown (*5, 9*) and knockout (*10*) studies, as well as by biochemical (*9, 11, 12*), microscopy (*4, 5, 13*), and proximity labeling (*14*) studies identifying ATG2 as an LD localized protein. Furthermore, ATG2A/B deficiency leads to LD accumulation and defects in fatty acid trafficking from LDs (*10*). LD biology is regulated by ATG2 independently from its canonical role in autophagosome biogenesis based on experiments demonstrating that disruption of other essential autophagy machinery does not lead to LD accumulation (*5, 10*). Finally, close apposition and membrane deformation between the phagophore and the LD can also be observed (*15*) suggesting the additional possibility that ATG2 might link these organelles. Whether ATG2-mediated lipid transport is involved in LD biology and indeed, even whether ATG2 is able to facilitate lipid transport from a phospholipid monolayer of an LD, remain unknown.

VPS13 family proteins intrinsically support two functions at MCSs: physically bridging two membranes to promote contact formation (*16*) and potentially transporting lipids between the membranes. Because of this, a knockout model cannot disentangle lipid transport activity from potential structural functions. Here, we demonstrate that ATG2A can transport lipids from phospholipid monolayer, and we test the hypothesis that ATG2-mediated maintenance of normal LD mass is dependent on ATG2’s lipid transport activity *in vivo*. We employ and expand on an approach originally used to study *in vivo* lipid transport by VPS13 which introduces multiple hydrophobic to hydrophilic mutations to the hydrophobic groove to obstruct lipid flow (*17*). We successfully generated a transport dead ATG2A mutant (TD-ATG2A) that maintains binding interactions with proteins and membranes bound by the WT protein and show that proper regulation of LD mass is dependent on the bridge-like lipid transport activity of ATG2A.

## Results

### ATG2A transports lipids from a phospholipid monolayer

ATG2 has been shown to facilitate lipid exchange between liposomes *in vitro (3, 6-8*). However, the phospholipid monolayer of an LD exhibits biophysical properties distinct from that of the phospholipid bilayer of liposomes or most cell membranes (*18, 19*). Therefore, to test whether ATG2A is capable of transporting lipids to or from a monolayer, we generated artificial lipid droplets (aLDs, also called adiposomes) (*20*) and used them as a substrate in lipid transfer assays. Artificial LDs are generated by differential centrifugation of a mixture of neutral triacylglycerol (TAG) and phospholipids, yielding a particle with a uniform phospholipid exterior and neutral lipid interior (*20*). Unlike liposome preparation, aLD preparation produces a heterogenous mixture of liposome-like particles and aLDs, which must be separated by differential centrifugation. aLDs displayed dark staining in positive stain electron microscopy, and could easily be differentiated from liposome-like particles (**Figure 1A**). Less than 2% of total particles identified in our aLD preparations displayed liposome-like morphology (**Figure 1A, S1A**). The median aLD particle was 165 nm in diameter with an inner quartile range from 129 nm to 211 nm (**S1A**). This is consistent with published aLD preparations (*20*) and allowed us to compare lipid transport from these particles against lipid transport on similarly sized liposome preparations.

**Figure 1.**
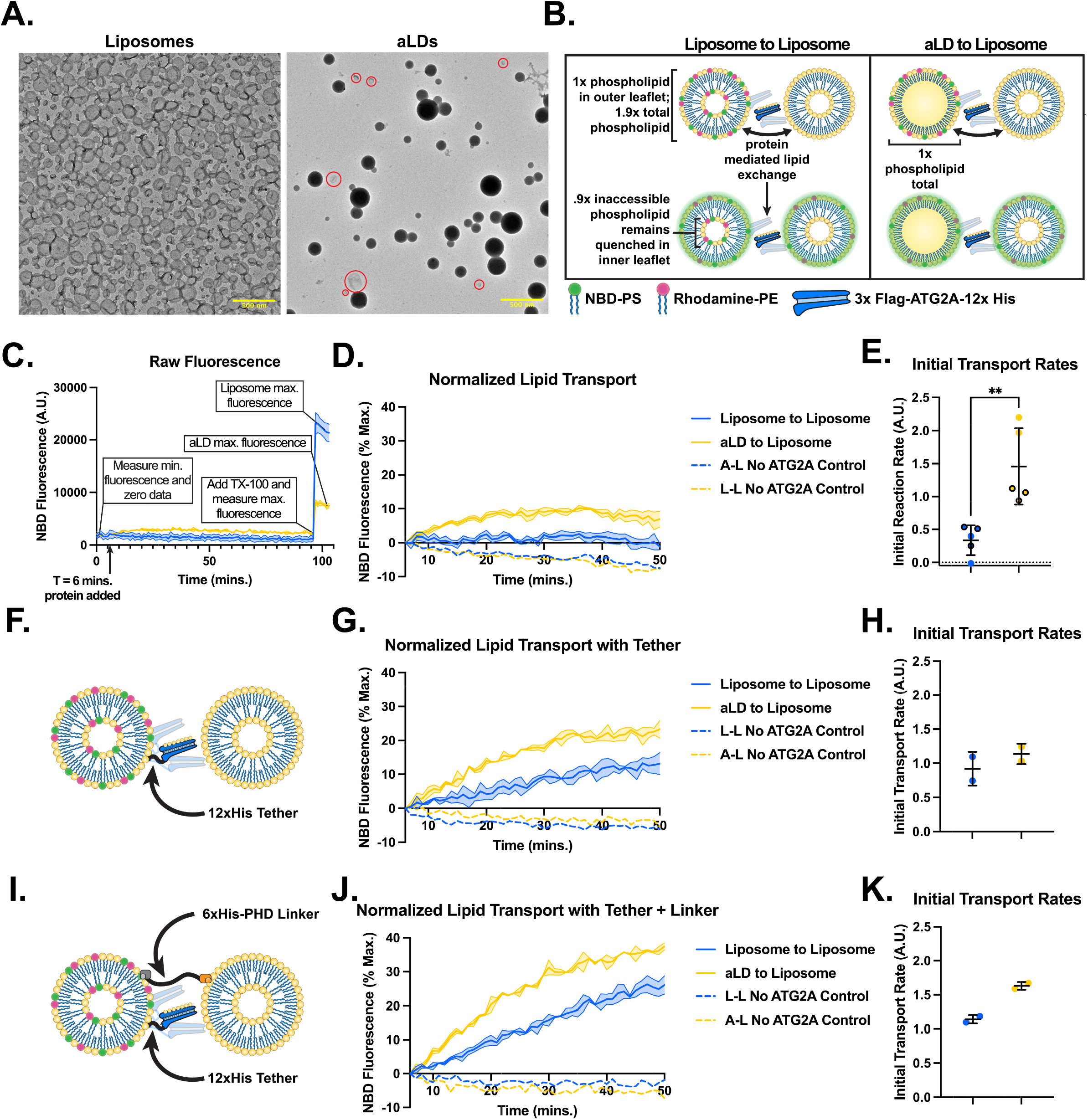
ATG2A transports lipids from phospholipid monolayers faster than phospholipid bilayers. **A)** Positive stain electron microscopy of donor liposomes and aLDs (91% DOPC, 2% rhodamine-PE, 2% NBD-PS, 5% DGS-NTA(Ni)). Liposome-like particles remaining in the aLD prep circled in red. **B)** Schematic of variations of FRET dequenching lipid transport assay. 0.525x total phospholipid of aLDs (calculated against standard curve of liposomes) is added to each reaction to reach equivalent outer leaflet phospholipid for each type of membrane. ATG2 is depicted shuttling between membranes but bridge-like lipid transport is also likely. **C)** Raw NBD fluorescence readings of FRET dequenching lipid transport assay. Protein added to assay at 6 minutes; reaction is fully dequenched with triton x-100 at the endpoint. **D)** Lipid transport between 0% PE donor membranes and 15% PE acceptor membranes. Reaction plotted as % maximum fluorescence. Fluorescence values before protein addition are set as 0% and fluorescence readings after triton x-100 addition are set as 100% fluorescence. Dashed reaction traces represent control reactions containing donor and acceptor liposomes with no protein. **E)** Initial reaction rates plotted from first 10 minutes of aLD to liposome and liposome to liposome assays. Reaction rates are normalized to donor + acceptor membrane control prior to quantification. Data plotted in D indicated by circled data points. ** p <0.01; Welch’s T-test. **F**,**G)** Reaction as in D, but donor membranes contain 5% DGS-NTA(Ni) to bind the COOH-terminal 12xHis tag on ATG2A. Acceptor membranes contain 5% PI(4,5)P_2_. **H)** Quantification of reaction rates identical in G. **I**,**J)** Reaction as in F,G, but an artificial construct containing a 6xHis tag, GGGGS linker, and a PI(4,5)P_2_ binding PH domain also included (6xHis-PHD Linker) to bring donor and acceptor membranes into close proximity. Dashed reaction traces represent donor + acceptor + linker controls. **K)** Quantification of reaction rates as in G.

To test whether ATG2A can transport lipids to and from aLDs, we used a FRET dequenching assay as in (*3*). In our system, donor liposomes or aLDs carry both 2% NBD-PS and 2% rhodamine-PE while acceptor membranes are initially without fluorophores. As lipids are exchanged, the surface density of the fluorescent lipids is reduced, rhodamine no longer fully quenches NBD, and NBD fluorescence can be detected (**Figure 1B**). Because the properties of monolayers and bilayers differ (*19*), we compared liposome-to-liposome lipid transport to aLD-to-liposome lipid transport (**Figure 1C**). In order to reveal the extent of lipid exchange, lipid-mixing assays are typically normalized to a fully-dequenched value achieved after the full solubilization of the lipids in detergent (**Figure 1C, S1B-C**); (*21*)). However, the mixing of liposome and aLD data poses a problem, because liposomes will harbor an additional membrane leaflet that does not participate in lipid exchange *a priori* but contributes to the detergent maximum (**Figure 1B-C, S1C**). Thus, we present our data with a normalized detergent maximum, in which bilayer values have been multiplied into a theoretical fluorescence maximum equivalent to a monolayer (approximately 1.9x the measured detergent maximum as determined by dithionite assay) (**Figure 1C,D, S1B-C**).

Addition of recombinant ATG2A (**Figure S1D i**) to the liposome-liposome reaction resulted in minimal lipid transport (**Figure 1D,E**). Note that, unlike previously published assays, we did not promote the recruitment of ATG2A to either membrane with artificial tethers (*3, 22*), adaptor proteins (*7, 8, 23*), or high phosphatidylethanolamine (PE) percentages. Thus, ATG2A alone is not an efficient lipid transporter on ∼100 nm liposomes with 0% PE. In stark contrast, addition of ATG2A to reactions where the donor membranes were aLDs of equivalent phospholipid composition led to a rapid and robust increase in fluorescence **(Figure 1E)**, indicating significant lipid exchange. Crucially, in all cases, mixing liposomes or aLDs alone did not lead to any appreciable change in NBD fluorescence (**Figure 1D, S1E; dashed lines**).

To establish a relative rate of lipid transport, we substracted the fluorescence values from protein-free controls (dashed lines in Figure 1D, which decline likely due to bleaching) from the protein-containing reactions. The inclusion of a monolayer structure accelerates lipid-mixing at least 4-fold over 100 nm liposomes of identical phospholipid composition (**Figure 1E**). Thus, ATG2A is an efficient and robust transporter of phospholipids residing in lipid monolayers.

We hypothesized that the transport rate differences between liposome-liposome and aLD-liposome reactions may be due to differential engagement of ATG2A with each membrane. Therefore we asked whether artificially tethering the C-terminus of ATG2A to the donor membrane by a 12xHis tag-DGS-NTA(Ni) lipid interaction could recover lipid transport from 0% PE liposomes (**Figure 1F**). The tether restored ATG2A-dependent lipid transport (**Figure 1G**) to approximately 80% of the initial rate of reactions where the donor membrane was substituted for an aLD (**Figure 1H**). Finally, a key challenge with studying lipid transfer on lipid droplets is that these structures can be driven to spontaneously coalesce much more easily than stable bilayers (*24*). All of our ATG2-free controls, including aLD-liposome controls, show no evidence of lipid exchange (see dashed lines in Figures 1D,G; S1E), however, we endeavored to stress the system by maximizing collisional frequency of these particles. Thus we repeated the experiment in Figure 1G, but also included a membrane-membrane linker (*3*) which forces all particles into relatively close apposition **(Figure 1I)**. Note that even in the presence of this linker, there is no evidence for lipid exchange unless ATG2 is present. (**Figure 1J-K; S1F**).

### ATG2A directly and strongly binds lipid droplets

Several groups have shown that recombinant ATG2A can localize to LDs in cells through its C-terminal amphipathic alpha helices (ATG2-C domain (**Figure 2A**)) (*4, 5*), and in fact, preferentially accumulates at those organelles. Whether that interaction is direct or depends upon other cellular factors has been unknown. Our observation of lipid transport in Figure 1 implies that ATG2A can bind directly to aLDs. Further, we hypothesized that faster lipid transfer in the aLD-based assay might reflect a more stable membrane engagement of ATG2A to monolayers over bilayers. We can test both questions directly with in vitro mimics of LDs.

**Figure 2.**
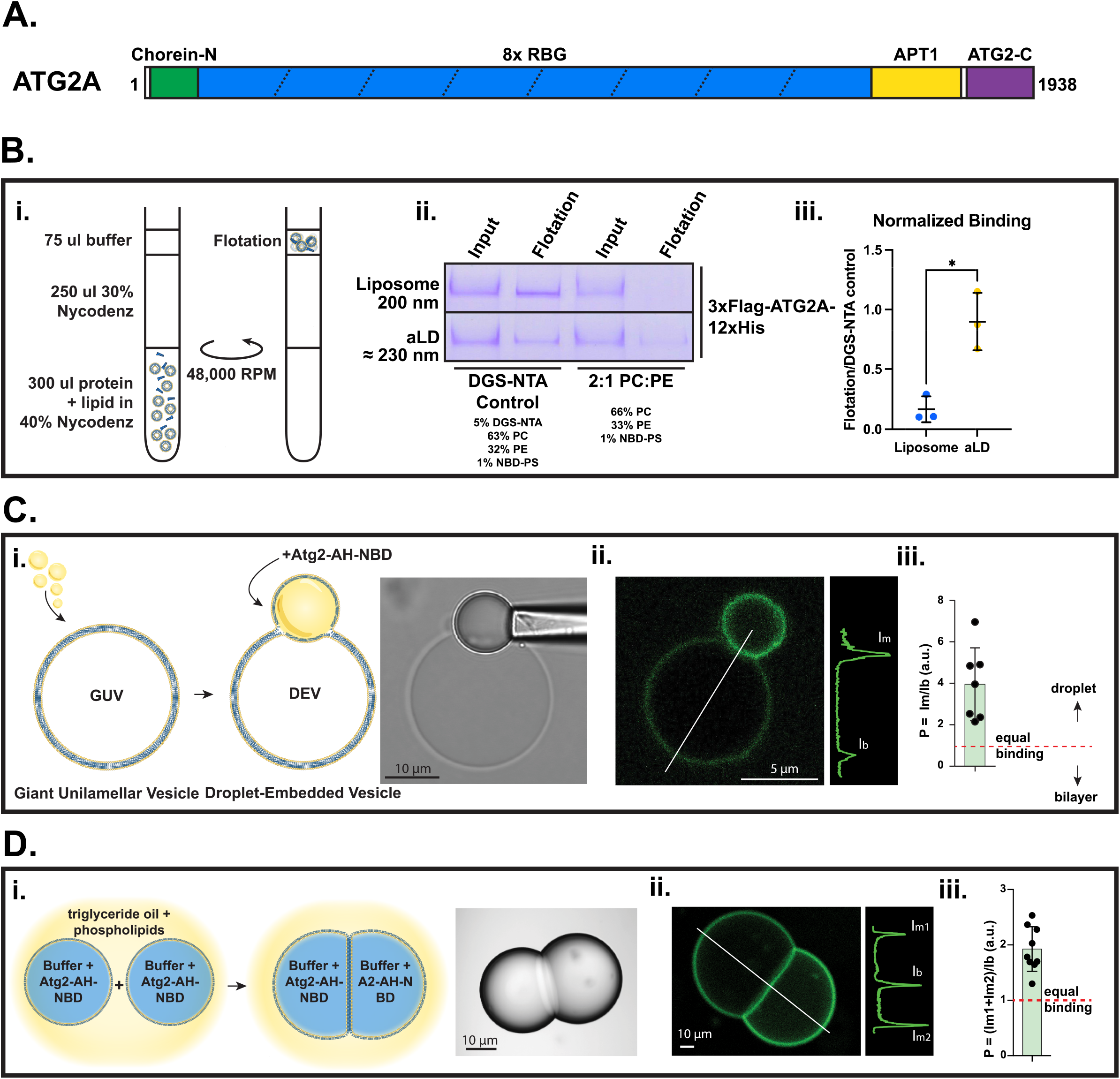
ATG2A binds lipid droplet surfaces directly via its amphipathic helical domain. **A)** Schematic of ATG2A domains. **Bi)** Schematic of membrane flotation assay. **Bii**) Coomassie stain of input samples and flotation samples from 3xFlag-ATG2A-12xHis flotation assay. Control membranes contain 5% DGS-NTA(Ni) and 2:1 DOPC:DOPE, and experimental membranes contain 2:1 DOPC:DOPE. 1% NBD-PS included to quantify aLDs. aLDs are loaded at .525x total phospholipid relative to liposomes, and the protein to outer leaflet phospholipid is 1:1000. 6.25% of input and flotation are loaded onto gel. 2:1 DOPC:DOPE aLDs are approximately 230 nm in diameter (Wang et al. (*20*)) so 200 nm extruded liposomes were compared. **Biii)** Quantification of liposome and aLD flotation assays with each sample normalized as ((flotation/input)/(DGS-NTA(Ni) flotation/ DGS-NTA(Ni) input)). Replicates represent three independent membrane preparations. * p <0.05; Welch’s T-test. **Ci)** Schematic of droplet-embedded vesicle construction. **Cii)** Representative fluorescent micrograph and linescan of NBD-Atg2-AH. **Ciii)** Quantification of NBD-Atg2-AH monolayer:bilayer ratio. **Di)** Schematic of the droplet interface bilayer assay. **Dii)** Representative fluorescent micrograph and linescan of droplet interface bilayers. **Diii)** Quantification of monolayer:bilayer ratio.

To test ATG2A membrane binding, we mixed 3xFlag-ATG2A-12xHis with liposomes or aLDs of approximately equivalent size and equivalent phospholipid (PL) composition and then floated these membranes on density gradients to look for ATG2A associating with floated materials (**Figure 2B i**). Based on published measurements, aLDs with 2:1 DOPC:DOPE ratios should be approximately 230 nm (*20*). We compared binding between these aLDs and liposomes extruded to 200 nm. Membranes with 5% DGS-NTA(Ni) that stably bind the C-terminal 12x histidine tag on ATG2A served as positive controls for establishing maximal binding (**Figure 2B ii**). When ATGA was mixed with 66%PC/33%PE liposomes, we only recovered 17% of the DGS-NTA(Ni) control. In contrast, when aLDs were used with the same phospholipid composition, ATG2A recovery in the floated fraction was 90% of the DGS-NTA(Ni) control (**Figure 2B iii; S2A**). Thus, even in this non-equilibrium binding assay, ATG2A stayed stably associated with aLDs throughout the float, with a final recovery approximately 5.5-fold higher than that of liposomes.

The binding of ATG2A to LDs in cells relies on an amphipathic alpha-helix located in a bundle of alpha-helices at its C-terminus (*4*). To investigate whether this motif exhibits a preference for associating with monolayers as opposed to bilayers, we produced a synthetic peptide from the yeast Atg2 protein that corresponds to this domain in human ATG2. To follow its localization, a fluorescent NBD-GG moiety was included at its N-terminus (NBD-GG-AH-Atg2). Equilibrium conditions were utilized to assess peptide binding to both monolayers and bilayers in two complementary *in vitro* membrane model systems.

In the first system, we generated giant unilamellar vesicles (GUVs) with a composition resembling the endoplasmic reticulum (ER), and subsequently mixed them with artificial triolein LDs to create droplet-embedded giant unilamellar vesicles (DEVs)(*25*). These DEVs featured an aLD embedded within the GUV bilayer (**Figure 2C, S2E**). The addition of NBD-GG-AH-Atg2 to DEVs demonstrated a preferential binding to the aLD, as indicated by the approximately 5-fold enrichment of NBD signal on the monolayer surfaces compared to the bilayer signal (**Figure 2Ciii**).

To further examine the preference for monolayers, we utilized a droplet interface bilayer setup, which allowed us to compare the binding preference of phospholipid monolayers versus bilayers (*26*). This system involved the creation of buffer-in-triglyceride droplets encapsulating the peptide, each covered with a phospholipid monolayer. Upon their proximity, the droplets spontaneously adhered to form a bilayer between the two droplet monolayers (*27*), similar to the DEV system. By analyzing the distribution of green signal to different surfaces, we calculated a binding coefficient to assess the monolayer/bilayer preference (**Figure 2D, S2E**). Consistent with the DEV experiment, AH-Atg2 displayed a marked inclination towards interacting with the monolayer surfaces.

Together, our results show that ATG2 proteins can bind directly to LD droplet surfaces, bind these surfaces with greater affinity than bilayers and that this preference is encoded in the membrane-interaction attributes of the COOH-terminal amphipathic helices.

### Bridge-like lipid transport is favored upon tight membrane binding

Depletion of ATG2 proteins in cells leads to dramatic changes in LD size and number (*5, 9, 10*), but whether these phenotypes depend upon the lipid-transport capacity of ATG2 has been uncertain. To test this question, we developed a lipid transport dead mutant protein. Our strategy is an extension of two previous studies: 1) we have shown that clusters of mutations within the amino-terminus of the hydrophobic channel can block lipid transfer by fragments of ATG2 (*3*), and we and others have shown that the introduction of clusters of mutations designed to prevent lipid migration through the channel (and thus blocking bridge-like transport) also block biologically relevant activities of the protein when introduced in cells (*17, 28*). Thus here, we introduce ten mutations into full-length ATG2A (**Figure 3A-B**) to prevent lipid movement down its hydrophobic core, generating a transport-dead ATG2A (TD-ATG2A). *In silico*, we tested potential clusters of mutations that would produce minimal structural perturbation, using the OmegaFold prediction program (*29*), settling on a combination predicted to fold similarly to WT ATG2A, with the secondary and tertiary structures of the protein mostly preserved except in the immediate proximity of the mutations (**Figure 3A-C**). Critically, like WT ATG2A, we were able to express and purify preparative quantities of TD-ATG2A that were sufficient for quality control structural analysis (**Figure S2F**) and subsequent *in vitro* lipid transport assays, suggesting comparable levels of protein stability. Negative stain electron microscopy of the purified mutant protein identified particles that sorted into 11 rod-like classes containing 6,300 of 18,000 total particles. Rod-like particles were approximately 20 nm in length, similar to published lengths of WT ATG2A (**Figure 3D**) (*3*) and to our own recombinant WT ATG2A. Thus, gross morphology is maintained, even as OmegaFold predicts that the clustering of mutations at the N-terminus will disrupt and occlude the hydrophobic channel near the chorein-N motif, potentially blocking lipid transport (**Figure 3C**).

**Figure 3.**
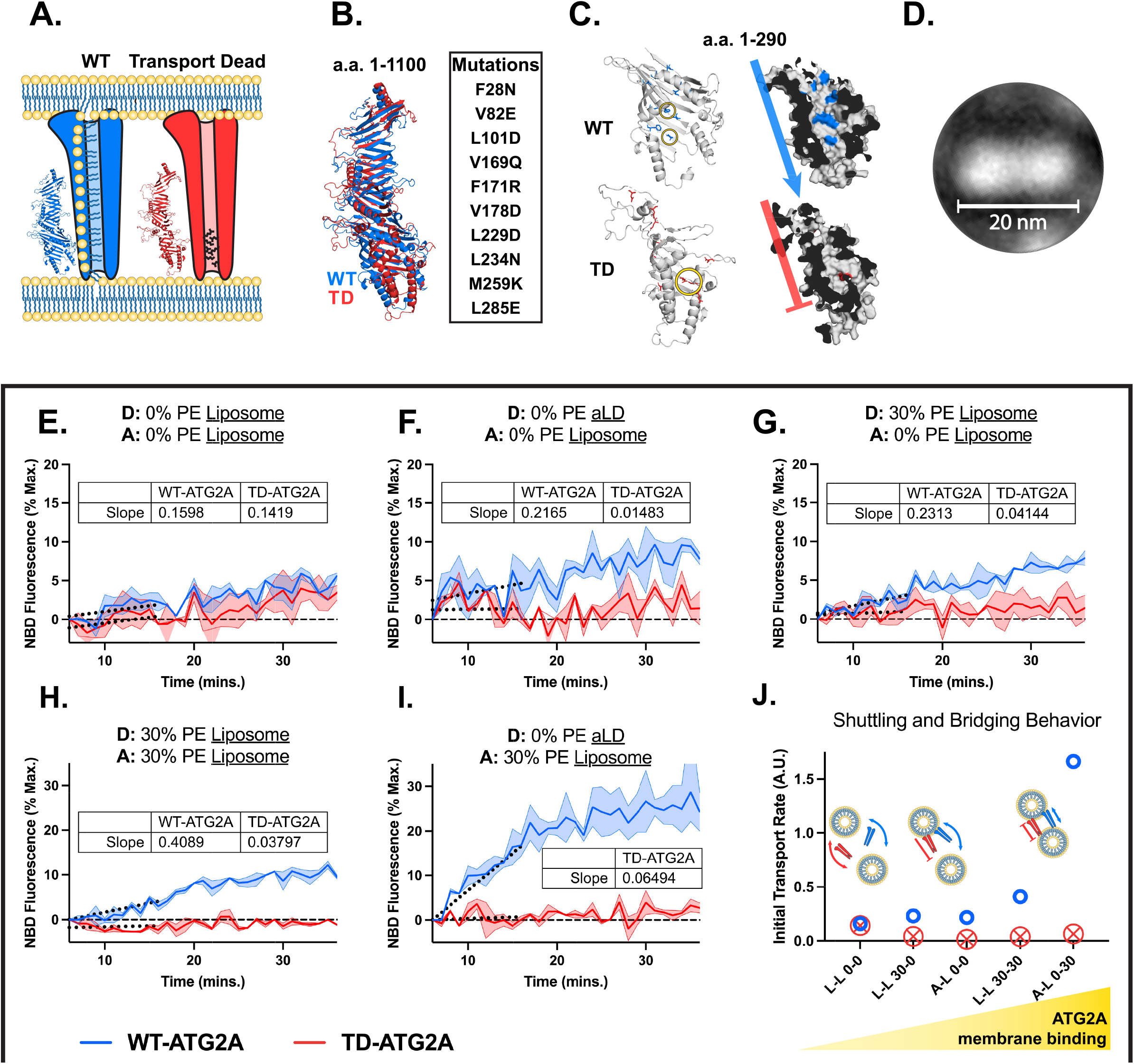
Mutation of hydrophobic channel dramatically impairs bridge-like lipid transport through ATG2A. **A)**. Schematic of mutation strategy employed to block lipid transport in ATG2A. Hydrophobic to hydrophilic mutated residues in the N-terminus of the transport dead mutant are depicted in black. **B)** OmegaFold structural predictions of the first 1100 amino acids of WT and TD ATG2A overlayed. Mutated residues are listed to the right. **C)** Omegafold structural predictions of the N-terminal 290 amino acids of WT and TD ATG2A. Surface renderings predict that the transport dead mutant hydrophobic channel is collapsed by interactions between two hydrophilic residues circled in yellow. Mutated residues are colored. **D)** Representative class from eleven rod-like classes (all shown in **Fig. S2**) identified by negative stain electron microscopy of purified 3xFlag-TD-ATG2A-12xHis. **E-I)** Lipid transport assays with 50 nM 3xFlag-WT-ATG2A-12xHis or 50 nM 3xFlag-TD-ATG2A-12xHis between donor membranes containing 2% rhodamine-PE, 2% NBD-PS, the indicated amount of DOPE, and the remainder DOPC. Reactions are plotted with the no ATG2A control subtracted out to remove the declines in fluorescence related to bleaching. Slope was calculated from linear region of curves. **J)** Plot of lipid transport rates from E-I with illustrations of the hypothesized molecular dynamics for each reaction.

To establish whether TD-ATG2A is actually transport-dead, we tested our recombinant protein *in vitro*. In an experimental system similar to Figure 1, in which membrane organelles are not tethered together and without affinity-lipids to specifically recruit his-tagged ATG2A to membranes, we detect a weak ATG2A-dependent lipid transport from liposome to liposome (**Figure 3E**). Intriguingly, the TD-ATG2A retains comparable activity. In contrast, when lipid transport between a liposome and an aLD is tested, we again observe substantial ATG2A-mediated lipid transport from aLDs to liposomes; however, we do not detect any lipid transport by TD-ATG2A (**Figure 3F**). Based on the membrane-surface binding results in Figure 2, the key difference in these two experimental systems is that with an aLD present, ATG2A maintains a prolonged interaction with the aLD surface while in the liposomes-only experiment, ATG2A is expected to predominantly be in solution. We suspect that on aLDs, only bridge-like transport along the length of ATG2A is possible and thus the mutations fully abrogate lipid exchange, while in the liposome experiment where ATG2A binds only weakly, ATG2A can function as a shuttle, potentially extracting and depositing lipids from just the unmutated, COOH-terminal end of the protein (**Figure S2H**).

Monolayers around LDs are replete with lipid-packing defects which are thought to promote the recruitment of proteins bearing amphipathic helices, like ATG2, to these organelles. We can mimic these defects on bilayers by introducing high concentrations of cone-shaped lipids such as phosphatidylethanolamine (PE) into our liposome system. Increasing PE percentages on either the donor, the acceptor or both membranes up to as much as 30%, drives more ATG2A membrane interaction and also drives a more efficient lipid transport **(Figures 3G-H, S2 B-D)**. However, TD-ATG2A is essentially inactive when 30% PE liposomes are included as either the donor or donor and acceptor membranes (**Figures 3G-H**). Furthermore, inclusion of 30% PE into an aLD drove the most robust lipid transport in our system (**Figure 3I**), but again in this context of tight membrane binding, TD-ATG2A was now inactive.

Lastly, we tested TD-ATG2 in a system that included both a tethered form of ATG2 to maximize membrane recruitment and a linker between membranes meant to recapitulate the organization of closely apposed membranes expected at a contact site (**Figure S2I**). In this optimized assay, WT-ATG2 drives lipid transport efficiently regardless of whether the donor is an aLD or a liposome presumably because ATG2A membrane binding is strong in both cases. However, even as lipid transport becomes more efficient with the WT-ATG2, TD-ATG2 remains fully inactive on aLDs and only just barely retains detectable activity on liposomes. Thus, the strategy of including mutations within the hydrophobic core completely disrupts lipid transport when ATG2A is expected to maintain prolonged interaction with the membrane, i.e., when the only form of lipid transport available is bridge-like transport through the channel itself.

### ATG2A lipid transport is required for LD homeostasis

Finally, we tested whether bridge-like lipid transport is necessary to support ATG2-mediated LD activity in cells. To study TD-ATG2A without the confounding effects of endogenous protein, we stably expressed GFP-tagged versions of either WT- or TD-ATG2A in ATG2A/B double knockout HEK293 cells (ATG2 DKO) that we described previously (*3*). Under these conditions, we could achieve equivalent expression levels between the WT and mutant protein (**Figure S3A**). Next, to assess whether lipid transport *per se* is being isolated as a variable, we tested whether GFP-TD-ATG2A localizes correctly and can still bind the same binding partners as WT ATG2A. When LDs were isolated from cellular lysates using a density separation technique (*30*), both GFP-TD-ATG2A and GFP-WT-ATG2A were enriched in the Plin3-positive, LD isolates (**Figure 4A**). Furthermore, Both GFP-WT-ATG2A and GFP-TD-ATG2A localized to Bodipy-stained lipid droplets (**Figure 4B**).

**Figure 4.**
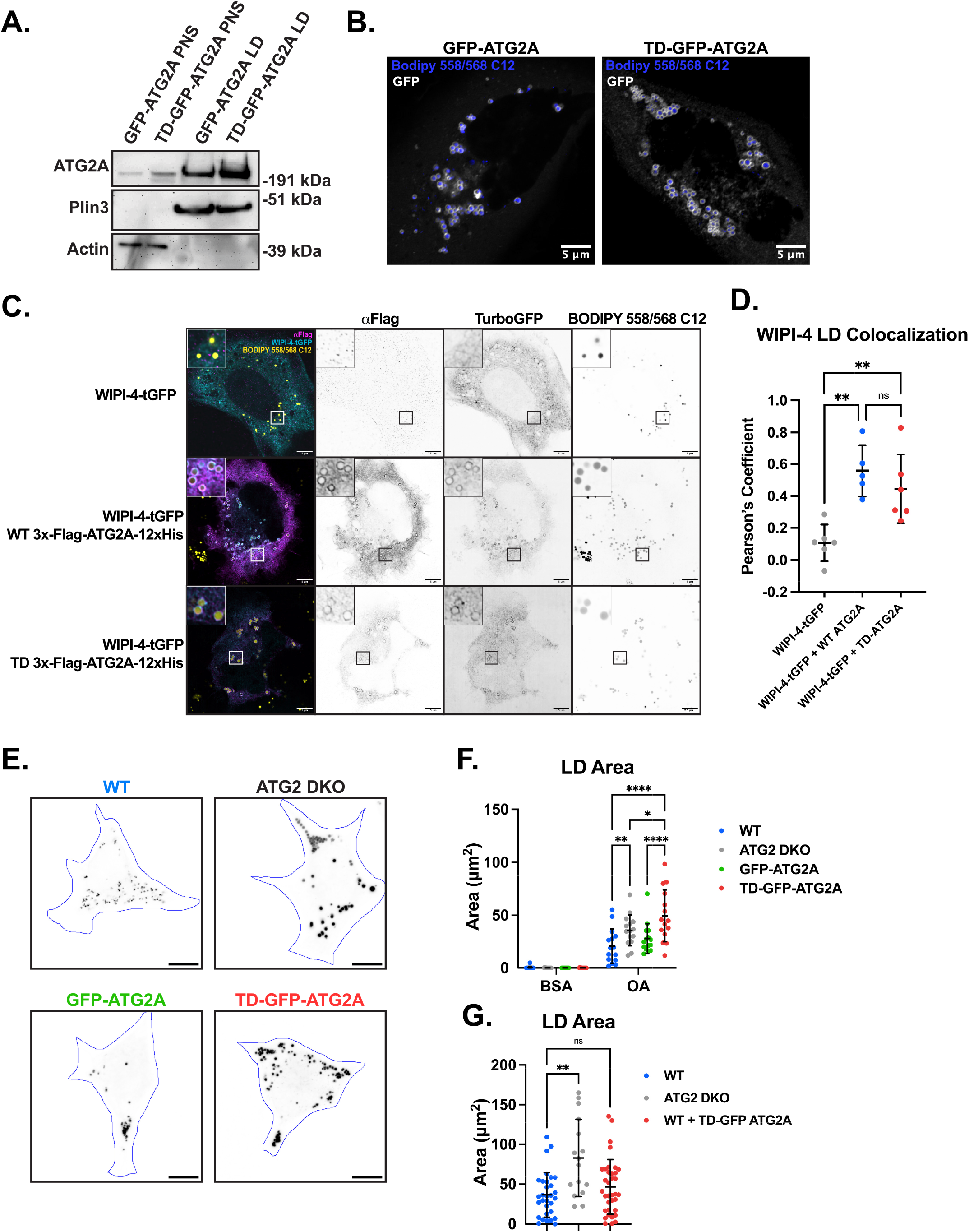
Loss of ATG2 mediated bridge-like lipid transport causes lipid droplet accumulation in cells. **A)** Immunoblot of post-nuclear supernatant fractions (PNS) and lipid droplet fractions (LD) from ATG2 DKO cells stably expressing either WT or TD GFP-ATG2A. Enrichment of LD is evidenced by PLIN3 intensity. **B)** Micrographs from live cell imaging of GFP-WT-ATG2A and GFP-TD-ATG2A transfected cells treated with 100 μM OA and stained with BODIPY 558/568 C12 (GFP in white and BODIPY in blue). Scale bars = 5 μm. **C)** Micrographs from fixed WT HeLa cells transfected with WIPI-4-TurboGFP alone or in combination with 3xFlag-WT/TD-ATG2A-12xHis. ATG2A is visualized by immunofluorescence against its N-terminal 3xFlag tag, and LDs are visualized by BODIPY 558/568 C12 staining. **D)** Quantification of colocalization between WIPI-4-tGFP and LDs. Pearson’s coefficients were calculated using FIJI colocalization threshold function. Each data point represents one transfected cell. ** p < 0.01; one-way ANOVA. **E)** Representative maximum projection micrographs of HEK293 cells treated with 100 μM OA and stained with BODIPY 558/568 C12. LDs are depicted in black and cell outlines are shown in blue. Outlines were generated manually after maximizing contrast. **F)** Quantification of total LD cross-sectional area from cells in E and corresponding BSA controls. * p < 0.05, ** p < 0.01, **** p < 0.0001; two-way ANOVA. **G)** Quantification of LD cross-sectional area from max projections of cells treated with OA. ** p < 0.01; one-way ANOVA.

The functionally-relevant binding partners of ATG2A on the LD (if any) are not yet known. On the autophagosome, ATG2 proteins bind to WIPI-4, via sequences near the COOH-terminus of ATG2 (*31, 32*). WIPI-4 is not generally considered an LD protein but can be recruited to LDs upon ATG2A overexpression (*33*). When we overexpress a TurboGFP-tagged form of WIPI-4, we do not observe significant accumulation on LDs in either WT HeLa cells (**Figure 4C**) or in our ATG2 DKO HEK293 cells (**Figure S3B**). Overexpression of a tagged WT ATG2A (3x-Flag-ATG2A-12xHis) leads to a dramatic redistribution of WIPI-4 onto LDs (**Figure 4C, S3B**) in both cell types. Critically, overexpression of the tagged TD-ATG2A drove a very similar level of redistribution, as assessed by Pearson’s correlation (**Figure 4D**). Moreover, WT and TD-GFP-ATG2A can Co-IP comparable levels of WIPI-4-TurboGFP (**Figure S3C**). We interpret these collected data to mean that TD-ATG2A is structurally similar to WT ATG2A, such that it retains its ability to traffic and localize to the same cellular compartments, and interact with the same proteins.

TD-ATG2A is, at a minimum, dramatically impaired for lipid transport *in vitro* (**Figure 3**). To test whether TD-ATG2A can support a lipid transport-dependent process *in vivo*, we measured macroautophagic flux in our ATG2 DKO cells with and without re-expression of ATG2 proteins. Autophagosome biogenesis requires the movement of millions of lipids to grow the autophagosome membrane (reviewed in (*34*)). The accumulation of the autophagosome marker LC3B following addition of bafilomycin can be used as a proxy for the total flux of membrane moving through the stress response pathway. As we have shown previously (*3*), ATG2 DKO cells do not flux LC3B at all, but instead maintain a very high constant level (**Figure S3A**). Re-expression of WT-GFP-ATG2 restores flux and also reverses the accumulation of an autophagy cargo receptor, p62. In contrast, TD-ATG2A cannot support autophagic flux in ATG2DKO cells (**Figure S3A**), even when expressed at levels many times over endogenous protein.

Whether ATG2-mediated lipid transport regulates LD biology is not known; thus we tested the functionality of TD-ATG2 in ATG2-dependent LD biology. Our HEK293 cells do not have easily detectable LDs under resting conditions, but feeding the cells OA for 4 hours leads to a strong accumulation of Bodipy-positive structures with an average cross-sectional area of about 25 μm^2^ (**Figure 4E,F**). As described by several other groups (*5, 9, 10*), genetic depletion of ATG2 leads to further LD accumulation. Stable re-expression of GFP-ATG2A returns total LD mass to WT-like levels. By contrast, expression of GFP-TD-ATG2A in ATG2 DKO cells did not rescue the LD phenotype (**Figure 4E,F**), again despite being expressed at levels significantly above endogenous protein. In fact, LD accumulation in TD-ATG2A rescued cells was consistently greater than in ATG2 DKO cells suggesting TD-ATG2A actually exacerbates the phenotype (**Figure 4F**). The mechanism of this enhanced phenotype is not yet clear; simple overexpression of TD-ATG2A in WT cells was without effect in LD biology or autophagy-ruling out a simple dominant negative model (**Figure 4E, S3C**). These results establish the lipid-transport capacity of ATG2A is key to LD biology.

## Discussion

Our work demonstrates lipid transport by ATG2A from a phospholipid monolayer, and establishes aLDs as a platform on which to explore lipid transport dynamics on phospholipid monolayers versus phospholipid bilayers. Furthermore, we present a framework for validating lipid transport-incompetent VPS13 family protein mutants, showing that structure, localization, and protein binding are preserved while lipid transport is blocked. Assessment of these parameters is essential to make claims about the necessity of lipid transport in a cellular process. We expect the approaches here will be generalizable to other members of the VPS13 family, several of which have also been implicated in LD biology (*22, 35, 36*).

Our findings establish that 1) ATG2 transports lipids more efficiently on a model phospholipid monolayer compared to a phospholipid bilayer, 2) ATG2 preferentially binds monolayers over bilayers, independent of other proteins, and 3) ATG2-mediated lipid transport is necessary for maintaining normal LD biology. Future efforts will be required to directly observe lipid transport to or from the LD in cells, and to determine the spatial organization of proteins involved in trafficking lipids at this membrane as well as the types of lipids transported (i.e. phospholipids, neutral lipids, fatty acids). To this end, recent work by Mailler et al. suggests the possibility of ATG2 mediated fatty acid transport (*10*).

In our in vitro assay system, it appears that the rate-limiting step involves the initial association of ATG2A with the membrane. Consequently, the more stable binding to aLDs, here promoted by higher packing defects, can explain a significant portion of the observed accelerated lipid transfer rate. Indeed, when considering monolayers, it’s important to note that phospholipids do not engage in trans-adhesive phospholipid-phospholipid interactions, unlike in bilayers (*19*). This implies that phospholipids are more readily available to access the ATG2 hydrophobic tunnel. Within the cellular context, a replenishment of phospholipids to the bilayer is required to sustain the lipid transfer, or an alternative is the reorganization of the bilayer’s integrity, potentially involving scramblase activity. All these considerations collectively contribute to an energetic deceleration of the transfer process from a bilayer compared to a droplet monolayer.

This phenomenon of packing defects exerting a comparable influence on ATG2 binding to aLDs and subsequent facilitation of lipid transfer also manifests on bilayers under specific conditions. *In vitro*, ATG2 lipid transport assays considering only bilayers have also reported accelerated lipid transfer when the bilayer structure or composition is modified to promote ATG2A recruitment, such as by inducing very high local curvature or introducing high phosphatidylserine concentrations (*6-8, 37*). On bilayers within cells, the major driver of this enhanced membrane affinity is probably other protein complexes, including WIPI family proteins or GABARAPs (*6-8, 23, 31*), while our work suggests that on monolayers, perhaps the natural affinity of ATG2A for LD surfaces is sufficient, because of the sparser phospholipid decoration (*19*) which is mimicked by having higher PE levels or inducing high curvature on a bilayer. As ATG2A may be constitutively associated with LDs, regulation of its lipid transport *in vivo* might ultimately depend more on the frequency of interaction with a second organelle, presumably the ER. Intriguingly, this framework would be consistent with findings by Cai et al. demonstrating by *in situ* cryo-EM tomography that VPS13C is frequently detached from its membrane binding partners (*16*). In light of these findings, our data are consistent with a model wherein a potential primary regulator of lipid transport among VPS13 family proteins is membrane binding.

## Acknowledgements

The authors would like to thank Shanta Nag for advising on *in vitro* experiments and other group members for helpful discussion. This work was supported by the National Institutes of Health to TJM (GM135290), KMR (R35GM131715), and DMF (F31AG079606). This work was supported by the Fondation pour la Recherche Médicale (FRM: EQU202103012564), the Programme Emergence de la Ville de Paris (DDEEES 165), and the ANR-16-TERC-0002-01 (LDEN) to ART. We thank the Yale Center for Cellular and Molecular Imaging Confocal Facility for usage of their Zeiss LSM 880 Airyscan confocal microscope. The authors would also like the thank the Center for Cellular and Molecular Imaging, Electron Microscopy Facility at Yale Medical School for assistance with the work presented here.

## Author Contributions

JLK and TJM conceived the study; JLK, TJM, ART, KMR designed experiments. JLK performed all *in vitro* biochemical experiments involving full length human ATG2A, all mammalian cell studies, and all statistical analysis. NW performed electron microscopy and assisted with protein purification. HE and LK performed droplet-embedded vesicle and droplet interface bilayer experiments with Atg2 peptide. MJP performed liposome flotation assays. DMF generated ATG2 DKO TD-GFP-ATG2A HEK293 cell line and cloned plasmid for full length WT/TD 3xFlag-ATG2A-12xHis and offered insightful conversation. TJM, ART, KMR oversaw all aspects of work performed in their respective labs. JLK and TJM wrote the manuscript with inputs from ART and KMR.

## Materials and Methods

### Materials

DOPE (Avanti Polar Lipids; 850725), DOPC (Avanti Polar Lipids; 850375), NBD-PS (Avanti Polar Lipids; 810198), Liss Rhod-PE (Avanti Polar Lipids; 810150), DGS-NTA(Ni) (Avanti Polar Lipids; 790404), PI_4,5_P2 (Avanti Polar Lipids; 840046), glyceryl trioleate (Sigma-Aldrich; T7140), BODIPY 558/568 C12 (Invitrogen; D3835), Imperial protein stain (Thermo Scientific; 24617), Flag M2 resin (Sigma-Aldrich; A2220), WIPI-4-TurboGFP (Origene; WDR45-tGFP transcript variant 1, RG209654), BSA-Oleate monounsaturated fatty acid complex and BSA control (Cayman Chemical; 29557 and 29556), rabbit anti-ATG2A (Cell Signalling; 15011), rabbit anti-ATG2B (Sigma-Aldrich; HPA019665), mouse anti-WIPI-4 (Santa Cruz Biotechnology; sc398272), rabbit anti-LC3B (D11) (Cell Signalling; 3868), rabbit anti-p62 (Cell Signalling; 95697), rabbit anti-Plin3 (Sigma-Aldrich; HPA006427), rabbit anti-beta-actin (Cell Signalling; 4970), mouse anti-Flag M2 (Millipore; F1804), rabbit anti-GFP (Cell Signalling; 2956), secondary antibodies, AlexaFluor 647 goat anti-mouse (Life Technologies; A21236), sodium dithionite Sigma-Aldrich (157953), DMEM (Gibco; 11965-092), MEM (Gibco; 31985-070), Expi293 medium (Gibco; A14351-01), 1x EBSS (Gibco; 24010-043).

### Liposome preparation

Liposomes were generated from lipids purchased from Avanti Polar Lipids. Lipids were combined according to molar ratios in borosilicate glass tubes then dried under nitrogen for 10-15 minutes followed by 1 hour drying in a vacuum chamber (*3*). Liposomes were resuspended in buffer A (500 mM NaCl, 20 mM HEPES, pH 8.0) by vortexing until dried lipid was no longer visible at the bottom of the tube. Resuspended liposomes were subjected to 7-10 freeze/thaw cycles to eliminate multilamellar structures, then extruded 21x to the appropriate diameter.

### aLD preparation

aLDs were generated by combining phospholipids according to molar ratios in borosilicate glass tubes. Glyceryltrioleate (TAG) was solubilized in chloroform and added to a separate tube from phospholipids in 5:2 (w/w) TAG to phospholipid ratio. All lipids were dried under nitrogen for 10-15 minutes followed by 1 hour drying in a vacuum chamber. TAG and buffer A were then combined in a single tube with dried phospholipids and aLDs were prepared according to Wang, et al (*20*). Substantial phospholipid is lost during differential centrifugation steps. Consequentially, the lipid concentration of the resulting aLDs is undefined. Lipid concentration must be known in FRET-based lipid transport assays, so phospholipid concentration of aLD preps was determined against a standard curve of fluorescent liposomes made from PLs in equivalent molar ratios to the aLDs. Fluorescence measurements were taken from membranes dissolved in a final concentration of 9% TritonX-100.

### aLD phospholipid quantification

Each aLD preparation was quantified by generating a standard curve of liposomes (1000 μM-62.5 μM PL). Liposome standards and aLD samples were dissolved in a final working concentration of 9% triton x-100 and 0.1x buffer A. NBD-fluorescence was then measured in a BioTeK Synergy H1 microplate reader.Total phospholipid content of aLDs was extrapolated from a trendline fit to the standard curve. Stock liposome phospholipid concentrations in subsequent assays were treated as 1.9-fold (0.525x) lower to account for inner leaflet lipids not participating in lipid transport or protein binding.

### Adherent cell culture

All cells were cultured at 37 °C, 5% CO_2_ in DMEM with 10% FBS and 1% penicillin/streptomycin. Cells were passaged by trypsinization.

### Expi293 culture and transfection

All proteins in this study were expressed and purified from Expi293 suspension cells. Cells were cultured at 37 °C, 8% CO_2_ in vented PETG flasks (Thermo Scientific, Nalgene; 4115-0500) with 125 rpm rotation. Prior to transfection, cells were grown to 3-5 × 10^6^ cells/mL then diluted to 2.5×10^6^ cells/mL and allowed to grow for 18-24 hours. Upon reaching 4.5-5.5 × 10^6^ cells/mL, the culture was diluted to 3 × 10^6^ cells/mL with fresh media. Cells were transfected with 100 μg plasmid per 100 mL of cultured cells. For every 100 mL of cells, 100 μg DNA was diluted in 5 mL OptiMEM. 300 µL of 1 mg/mL polyethylenimine linear 25k (Polyscience, 23966) was diluted into 5 mL OptiMEM in a separate tube. Both tubes were incubated for 5 minutes at room temperature then combined and further incubated for 30 minutes. The DNA/PEI mixture was then added dropwise into Expi293 cultures. 12-18 hours after transfection, protein expression was enhanced using 700 μL of 500 mM valproic acid. Cells were then harvested after 48-72 hours.

### ATG2A purification

Cell pellets were resuspended in 10 mL purification buffer A (500 mM NaCl, 20 mM HEPES, pH: 8.0, 10% glycerol, 1 mM TCEP) for every 100 mL of original culture volume. Purification buffer A was supplemented with 1x EDTA-free protease inhibitor cocktail (Roche; 11873580001). Cells were then lysed using one of two methods: Cell suspensions were subject to 6 freeze/thaw cycles in liquid nitrogen or lysed by nitrogen cavitation. Nitrogen cavitation was performed using a Parr Instruments 4369 cell disruption vessel pressurized to 500 PSI for 5 minutes on ice. Nitrogen bubbles were allowed to settle out of lysate. Lysates generated from either freeze/thaw cycles or nitrogen cavitation were then clarified by centrifugation at 18,500 x G, and supernatants were transferred to 150 μL of anti-Flag M2 agarose resin per 100 mL of initial culture volume (prewashed with 10 mL purification buffer A). Lysates were incubated with beads under mild agitation for 2 hours at 4 °C. Lysates and beads were then passed through a gravity filtration column and washed with 4 CVs of purification buffer. Beads were then incubated with 2.5 mM ATP and 5 mM Mg^2+^ in purification buffer overnight (WT ATG2A) or for 1 hour (TD ATG2A) at 4 °C. Columns were then washed with 2 CVs of purification buffer, followed by elution with 50 μL of 1 mg/mL 3x-Flag (DYKDDDDK) peptide (ApexBio Technology; A6001) per 100 mL of initial culture volume. Protein concentration was determined against a BSA standard curve by Coomassie staining prior to subsequent assays. Proteins were flash frozen in liquid nitrogen for storage at -80 °C. Protocol was modified from (*3*) to optimize recovery of TD-ATG2A.

### Atg2 Amphipathic Helix preparation

NBD-Atg2 amphipathic helix was chemically synthesized and purified by reverse phase high-performance liquid chromatography (HPLC) by Genic Bio. The purity was 95.17%, as determined by analytical HPLC and their mass was confirmed by mass spectrometry. The amino-acid sequence of the peptide is: NBD-GG RSFMAIGSGVKTLVTVLMSEYRQ (*S. cerevisiae* Atg2 amino acids 1350-1372). The peptide was solubilized in dimethyl sulfoxide (DMSO) and in HKM buffer (50 mM HEPES, 120 mM CH_3_CO_2_K, 1 mM MgCl_2_), with a final DMSO concentration of around 1.5%. The stock solution of the peptide was at 90 mM.

### Droplet-embedded vesicle assay

Phospholipids (DOPC/DOPE/cy5-DOPE (7:2.9:0.1) purchased from Avanti Polar Lipids Inc. were in chloroform (Sigma Aldrich) at 2.5 mM. They were spread on an indium tin oxide (ITO)-coated glass plate. After chloroform evaporation, the resulting lipid film was then placed under vacuum for 1 hour. The chamber was sealed with another ITO-coated glass plate. Giant unilamellar vesicles (GUVs) were grown by electroformation in a sucrose solution (0.1 g mL−1, ≈280 mosmol kg−1) with the following settings: 100 Hz, 1.55 V, for 2 hours. They were then collected carefully with a Pasteur pipette, placed in an Eppendorf® tube, and stored at 4 °C. The aLDs were made by mixing 5 µL of triolein and 100 µL of HKM. The mix was then vortexed and sonicated for 2 minutes to form droplets. DEVs were made by mixing GUV and aLD (1:1) and letting it turn on a spinning wheel for 15 min. A glass coverslip was pretreated with 10% (w/w) BSA and washed three times with buffer. 50 µL of HKM, 50 µL of DEV and 4 µL of NBD-Atg2 peptide (c=89,6 mM, 1.5% DMSO) were put on the glass coverslip and it was then observed by confocal fluorescence microscopy (LSM 800, Carl Zeiss, Oberkochen, Germany), with an oil-immersed ×63 objective.

### Droplet interface bilayer assay

The droplet-interface bilayers (DIBs) assay was performed following (*26*). Briefly, phospholipids (containing 0.1% of Cy5-PE) were evaporated under a stream of argon to remove the chloroform. The resulting lipid film was then resolubilized to the desired concentration (0.2% w/w) in trioctanoate/squalene (80/20),strongly vortexed and sonicated for 5 minutes to ensure complete mixing. To form droplet-interface bilayers (DIBs), buffer-in-oil emulsion droplets were made using 10 μl HKM buffer (50 mM HEPES, 120 mM potassium acetate, 1mM MgCl_2_ at pH 7.4) containing the peptide (1-3 mM) dispersed in 100 μL of the prepared oil solution containing the phospholipids. This mixture was vortexed in order to generate the emulsion droplets. The emulsion was then placed on a hydrophobic coverslip (glass coverslip #0 from Menzel Glaser, Braunschweig, Germany, which was covered by a reticulated polydimethylsiloxane film, deposited by spin coating). The sample was left to equilibrate for 10 minutes and was then observed by confocal fluorescence microscopy (LSM 800, Carl Zeiss, Oberkochen, Germany), with a ×10 or oil-immersed ×63 objective depending of the size of the droplets.

### Negative stain electron microscopy

Purified ATG2A was applied to glow discharged carbon film coated grids (CF400-CU; Electron Microscopy Sciences) for 1 minute and blotted for 2% uranyl acetate staining. The micrographs were taken on 120kV T12 at Yale CCMI facility, with a pixel size of 1.7A/pixel with a defocus of 1-1.5 μm. The micrographs were processed using Relion3.0.8 software: protein particles were manually picked after CTF correction for initial 2D classification, followed by autopicking and another round of 2D averaging (mask diameter: 280A).

### Positive stain electron microscopy

Performed as previously described (*20*). Briefly, 8 μL sample was loaded onto the glow-discharged carbon film coated grids (CF400-CU; Electron Microscopy Sciences) for 1 minute followed by blotting. The sample was then fixed with 1% osmium tetroxide for 10 minutes and washed three times with ddH_2_O. The sample was stained with 0.1% tannic acid for 5 minutes and 2% uranyl acetate for 5 minutes and washed with ddH_2_O for three time. The micrographs were acquired at 120 kV T12 with a pixel size of 0.75 nm at Yale CCMI facility.

### FRET dequenching lipid transport assay

Lipids were added to reaction with 25 μM total phospholipid donor membrane and 50 μM total phospholipid acceptor membrane. For reactions involving aLDs, they were added at 12.5 μM donor and 25 μM phospholipid to account for approximately half of liposome lipids being unable to participate in lipid transport. Each assay included a buffer only control, a donor + acceptor membrane only control for spontaneous fusion, a donor + acceptor membrane + triton x-100 control for initial maximum fluorescence, and a donor + acceptor + linker control for fusion when applicable. Fluorescence measurements were made using the BioTeK Synergy H1 microplate reader at 30 °C. Baseline fluorescence is measured for the first 5 minutes followed by protein or protein purification buffer addition to appropriate wells. Lipid transport reactions are measured for at least 30 minutes following protein addition, then 5 μL of 10% triton x-100 is added to each reaction and mixed to assess maximum possible fluorescence for each well. Reactions are quantified as percentage completion between pre-protein measurements and triton x-100 maximum fluorescence. Because liposome donor membrane reactions contain approximately 1.9x more total phospholipids (outer + inner leaflet), and consequently have a 1.9x higher detergent maximum for quantification, liposome percentage values are corrected by this factor to be compared to aLD donor membrane reactions.

### Membrane flotation assay

Protein and membrane were combined in a molar ratio of 1 protein to 1000 outer leaflet lipids. Binding reactions were performed in buffer A containing 2x protease inhibitor (Roche) for 1 hour at 4 °C. An input fraction was then removed and immediatedly combined in a 1:1 ratio with 4x LDS. The remaining reaction was combined 1:1 with 80% Nycodenz in buffer A (approximately 300 μL final volume) and loaded into the bottom of a Beckman 5 × 41 mm Ultra-clear ultra centrifuge tube (344090). 250-300 μL of 30% Nycodenz containing 1x protease inhibitor in buffer A was layered on top of the protein containing fraction. 75 μL of buffer A containing 1x protease inhibitor was applied to the top of the gradient. Gradients were then centrifuged at 48,000 RPM in a Beckmann SW55 swinging bucket rotor for 4-20 hours. Gradients were immediately frozen at -80 °C, and then flotation fractions were recovered by cutting the top of the ultracentrifuge tube containing 80-130 μL of the frozen gradient. Flotation fractions were thawed on ice and the volume was measured using a pipette. Flotation fractions were immediately combined in a 1:1 ratio with 4x LDS. Equal percentages of input fractions and flotation fractions were subject to SDS-PAGE with flotation percentages being corrected by a factor equivalent to (reaction volume)/(reaction volume – input volume). Protein was visualized by Coomassie staining and images were quantified by densitometry in FIJI.

### OmegaFold structural predictions

Structural predictions were performed using OmegaFold Colaboratory tool (*29*): https://colab.research.google.com/github/sokrypton/ColabFold/blob/main/beta/omegafold.ipynb

### GFP-(WT/TD)-ATG2A transient transfections

Transient transfections were performed with lipofectamine 3000 according to manufacturer protocols.

### Lipid droplet purification

Lipid droplets were purified from HEK293 cells after overnight treatment with 100 μM oleic acid BSA conjugate. Purification was performed according to (*30*). Lipid droplet fractions were harvested from the sucrose gradient by freezing the tubes and removing the top layer by cutting the tube with a razor blade. Equal volumes of LD suspension and 10% SDS in water were combined and incubated in a sonicating waterbath at 60 °C for 1 hour, vortexing the sample every 10 minutes. Samples were then centrifuged at 20,000 x G and the infranatant containing solubilized LD proteins was removed using a gel loading pipet.

### Immunofluorescence staining and colocalization analysis

Cells were grown at low density on glass coverslips in 24 well plates. 48 hours after transfection, cells were fixed by addition 500 μL of 4% paraformaldehyde in PBS was added to 1 mL of the culture media. After 5 minute incubation, culture media was replaced entirely with 250 μL of 4% paraformaldehyde in PBS, and cells were incubated at room temperature for 15 minutes. Each well was then washed 3x 5 minutes with 1x PBS at room temperature followed by blocking in 3% BSA, 0.1% saponin in PBS for 15 minutes. Coverslips were removed from culture plates and incubated face down with primary antibody in blocking buffer overnight at 4 °C in a humidified chamber. Coverslips were subsequently washed again and incubated with AlexaFluor conjugated antibodies at 1:600 dilution for 1 hour. After a final wash in PBS, coverslips were mounted to slides using Fluoromount-G (SouthernBiotech) and allowed to dry overnight before imaging. Colocalization was measured using ImageJ FIJI colocalization threshold tool. Pearson’s correlations are reported.

### Autophagic flux analysis

HEK293 cells were cultured in 6-well plates to 70-90% confluence. At the beginning of the assay, cells were washed with 1x PBS, and the media was replaced with fresh DMEM, DMEM containing 100 nM bafilomycin A1, or EBSS containing 100 nM bafilomycin A1. After a 2 hour incubation, cells were harvested by trypsinization and washed with PBS. Cell lysis was performed in RIPA buffer (50 mM Tris pH:7.4, 150 mM NaCl, 1% NP-40 substitute (v/v), 0.5% sodium deoxycholate (w/v), 0.1% sodium dodecylsulfate (w/v)) supplemented with 1x protease inhibitor cocktail (Roche). Lysates were clarified by centrifugation at 20,000 x G for 10 minutes, and protein concentration of clarified lysate was assayed by Pierce BCA protein assay (Thermo Scientific; 23225). 20 μg of protein from each sample were analyzed by SDS-PAGE and immunoblotting for LC3B and p62.

### Coimmunoprecipitation assay

WT HEK293 cells ATG2 DKO HEK293 cells stably expressing GFP-ATG2A or TD-GFP-ATG2A were transfected with WIPI-4-TurboGFP. WT cells were also transfected with a GFP construct as control for nonspecific binding. Cells once cells reached 100% confluence in 15 cm plates (48-72 hours after transfection), plates were harvested by cell scraping in ice cold PBS. Cell pellets were lysed by 10 minute incubation in 500 μL of AT2A Co-IP lysis buffer (10 mM Tris pH: 7.5, 150 mM NaCl, 0.5 mM EDTA, 0.5% (v/v) NP40 substitute (Roche; 11754599001)) supplemented with 1x EDTA-free protease inhibitor cocktail (Roche; 11873580001). Lysates were clarified by centrifugation at 20,000 x G for 10 minutes. 25 μL of GFP-Trap magnetic agarose affinity bead suspension was washed 3x in 1 mL of ATG2A Co-IP lysis buffer. 500 μL of clarified lysate was added to the beads and incubated at 4 °C with light agitation for 2 hours. Remaining lysate was reserved as input. Flowthrough was then discarded, and beads were washed 4x in 1 mL of Co-IP lysis buffer supplemented with 1x protease inhibitor cocktail. Protein was eluted from beads in 60 μL of 4x LDS sample buffer (Invitrogen; 2399548) diluted to 2x in ATG2A Co-IP lysis buffer. Samples were boiled for 5 minutes, and sample was removed from beads for SDS-PAGE.

### Lipid droplet area quantification

Lipid droplet cross sectional area was assessed from live-cell micrographs of HEK293 cells treated for 4 hours with 100 μM OA or BSA control. Cells were treated with BODIPY 558/568 C12 1:2000 in DMEM for 30 minutes prior to imaging. Images were captured using either the Zeiss LSM 880 Airyscan or the Nikon CSU-W1 SoRa microscope. Maximum projections were generated from each image, and minimum thresholds were set to exclude signal from BSA control cells. Cell outlines were manually established by maximizing contrast from background. Total area occupied by BODIPY signal within each cell was plotted.

### Statistical Analysis

Experiments with two groups were analyzed using Welch’s T-test to account for differences in variance. Experiments with three or more experimental groups were anaylzed by one-way analysis of variance (ANOVA). Experiments with multiple variables were analyzed by two-way analysis of variance. ANOVA was followed by a Holm-Sidak *post hoc* correction for multiple comparisons. Statistical analysis and data visualization were performed using Graphad Prism.

## Figure Legends

**Supplementary Figure 1.**
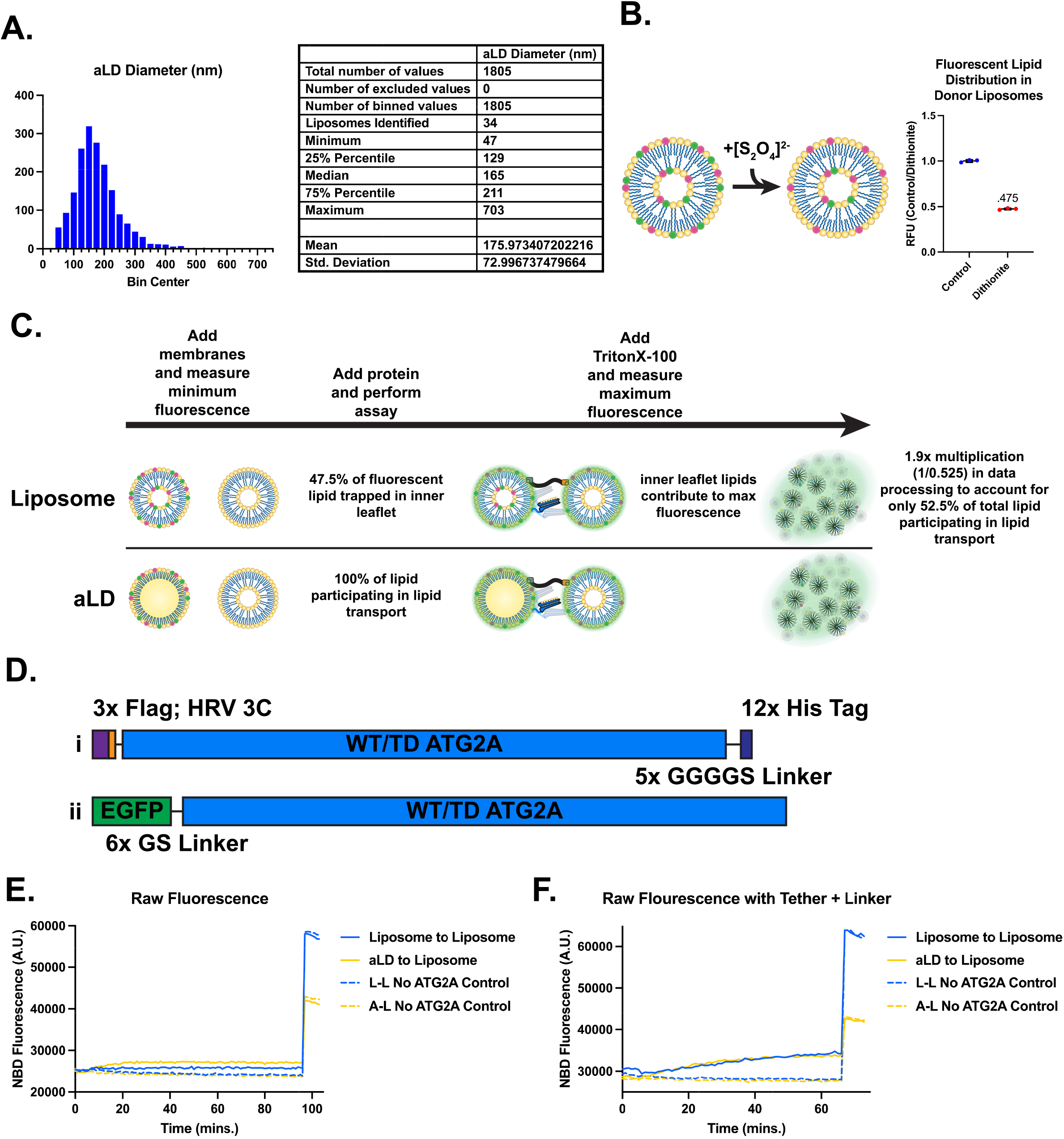
Characterization of in vitro liposome and aLDs and rationale for assay design. **A)**. Statistics on aLD size. aLDs were measured by identifying black particles in positive stain EM fields. Adjacent particles were segmented then Feret’s diameter was measured for each particle in FIJI. Liposome-like particles were identified manually. 34 of 1839 particles (1.8%) were liposome-like. **B)** Schematic of dithionite assay demonstrating quenching of external NBD on bilayer structures and ratio of remaining inner leaflet NBD fluorescent signal to total NBD fluorescent signal. **C)** Timeline of lipid transport assay and rationale for 1.9x correction to liposome-liposome transport values. **D)** Schematic of ATG2A constructs used in this study. **E)** Raw fluorescence readings from Figure 1D. **F)** Raw fluorescence readings from Figure 1J.

**Supplementary Figure 2.**
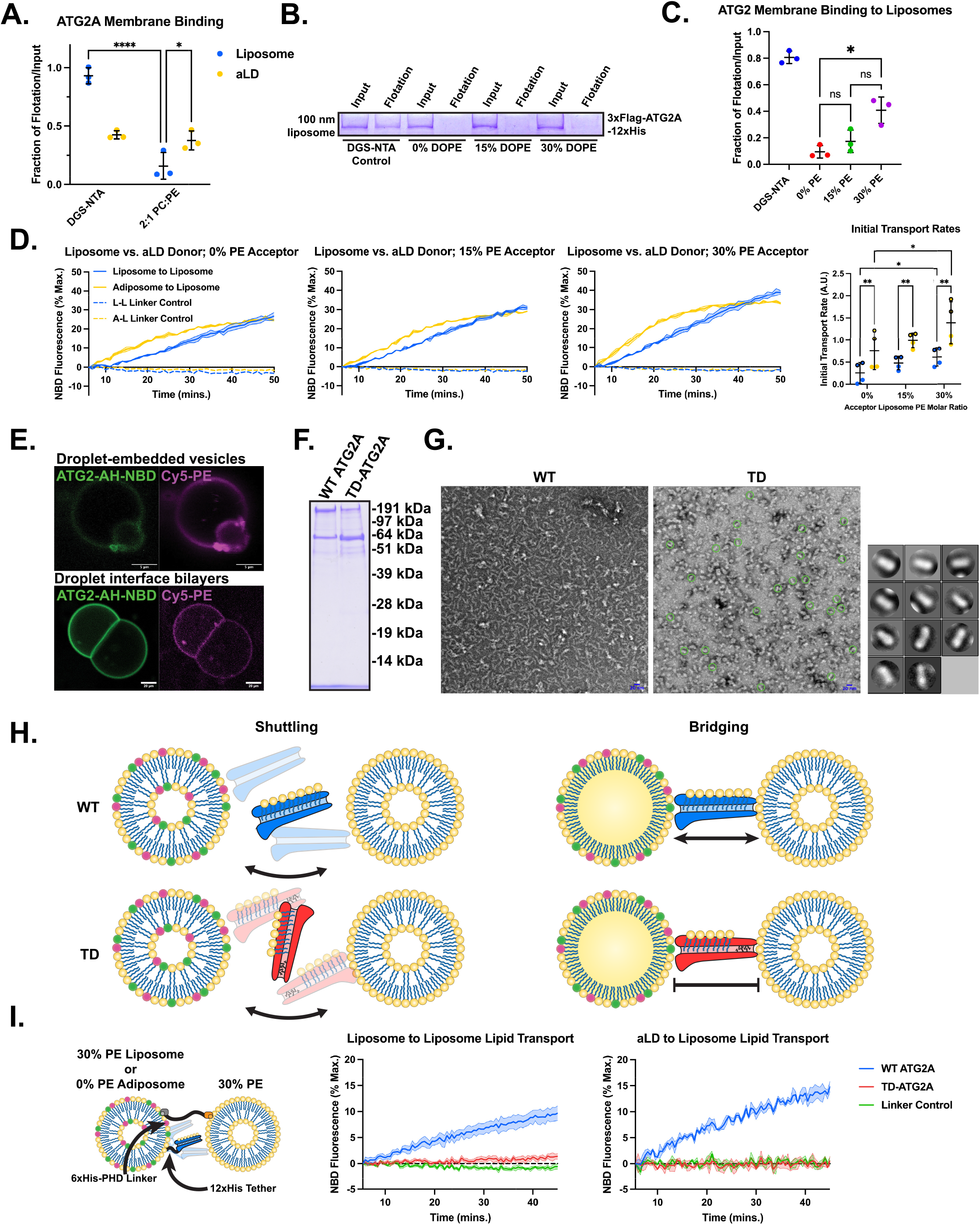
Membrane binding correlates with lipid transport rate. **A)** Raw flotation/input ratios for figure 2Biii. * p < 0.05, **** p < 0.0001; two-way ANOVA. **B-D** *<u>-- Membrane binding correlates with lipid transport rates</u>*. To directly explore the impact of membrane binding on lipid transport, we varied DOPE concentration in 100 nm liposomes representing acceptor membranes in the lipid transport assay to determine whether the introduction of PE mediated membrane packing defects could increase binding and accelerate lipid exchange. **B)** Coomassie stain of flotation assay performed identically to figure 2Bii, but with 100 nm liposomes representing acceptor liposomes in lipid transport assays. **C)** Quantification of flotation assays from three independent membrane preparations. * p < 0.05; one-way ANOVA. As hypothesized, we observed increases in ATG2A binding with 30% PE liposomes. **D)** Liposome to liposome and liposome to aLD lipid transport assays with 0% DOPE donor membranes and variable DOPE acceptor membranes as annotated. Reactions performed with 6xHis-PH linker construct and donor membranes containing 5% DGS-NTA(Ni) and acceptor membranes containing 5% PI(4,5)P_2_. Quantification of reaction rates from the first 10 minutes of each reaction. Data points represent technical replicates from two independent experiments. Data plotted in E annotated by circled data points. * p < 0.05, ** p < 0.01; two-way ANOVA. When membranes of increasing PE percentage are introduced as acceptor membranes to a lipid transport assay wherein the COOH-terminus of ATG2A is tethered to the donor membrane, we observe increases in lipid tranfer rate corresponding to the increases in ATG2A membrane binding to PE rich liposomes, suggesting that acceptor membrane PE promotes efficient lipid transport even when interacting with the N-terminus of ATG2A. Taken together, these data suggest that stable binding of ATG2A with each membrane participating in lipid transport is a primary determinant of transport rate *in vitro*. **E)** Representative images from monolayer distribution experiments comparing Atg-AH-NBD to phospholipid on each structure. **F)** Coomassie stain of 3xFlag-WT/TD-ATG2A-12xHis purification. **G)** Negative stain EM fields of WT and TD ATG2A from A. Rod-like particles are identified by green circles. 11 rod-like classes are shown which represent 33% of particles identified. **H)** Schematic of conceptual “shuttling” vs “bridge-like” transport modalities. **I)** lipid transport assays using a linker construct and DGS-NTA(Ni) to tether ATG2A to the donor membrane. Liposome-liposome reaction includes 3xFlag-WT-ATG2A-12xHis or 3xFlag-TD-ATG2A-12xHis between donor aLDs containing 0% DOPE, 2% rhodamine-PE, 2% NBD-PS, 5% DGS-NTA(Ni), and 91% DOPC and acceptor membranes containing 15% DOPE, 5% PI(4,5)P_2_, and 80% DOPC. Adiposome-liposome reaction includes donor liposome containing 30% DOPE, 2% rhodamine-PE, 2% NBD-PS, 5% DGS-NTA(Ni), and 61% DOPC. 30 nM protein in liposome assay and 15 nM protein in aLD assay. Data represent 3-5 replicates derived from at least 2 independent protein purifications and two independent liposome preparations.

**Supplementary Figure 3.**
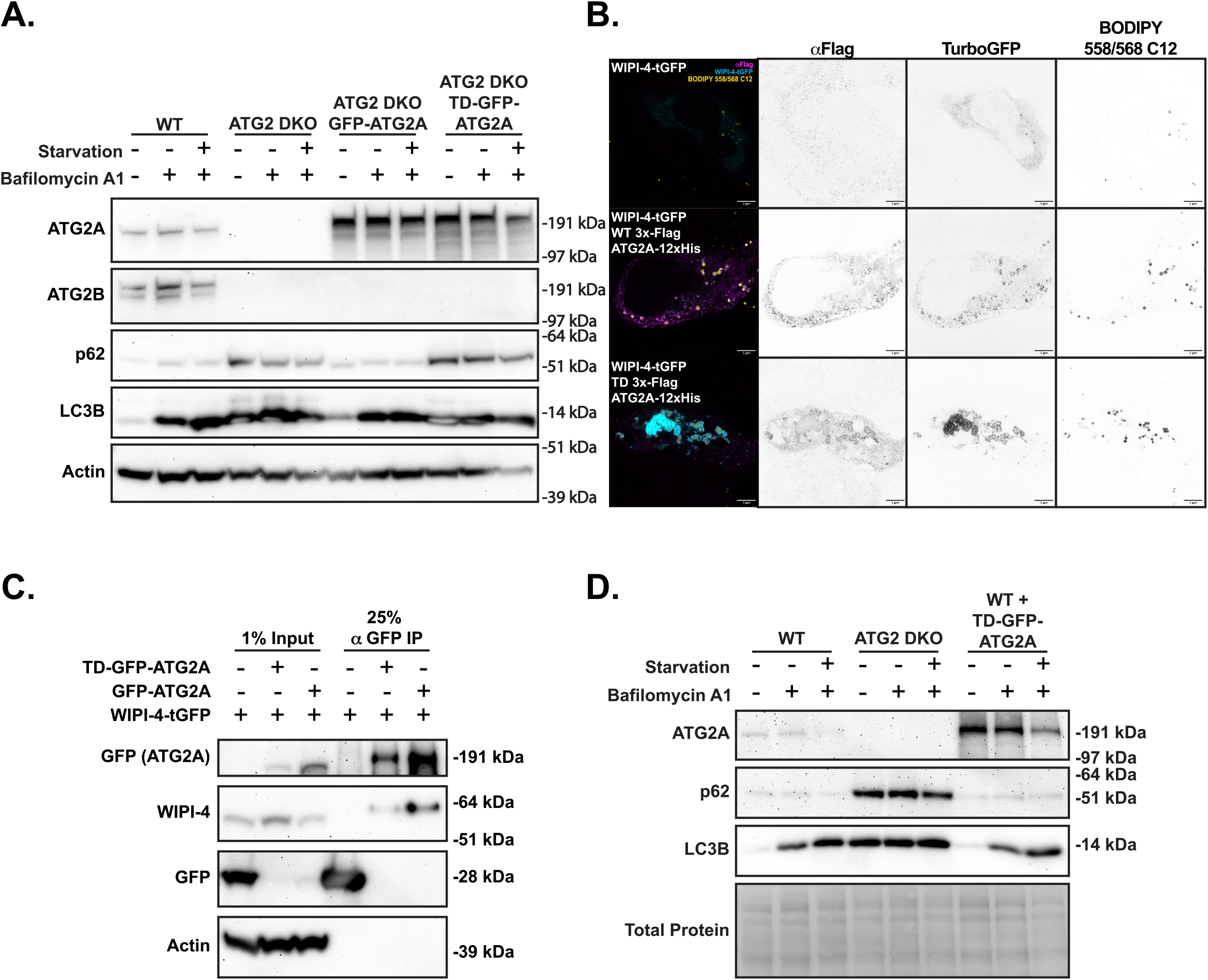
Further characterization of TD-ATG2A in cells. **A)** Autophagy is not rescued by expression of TD-ATG2A in ATG2A double knockout cells. Flux of an autophagosome marker (LC3) and flux of an autophagy cargo (p62) were followed in indicated HEK293 cells. Cells were treated with 100 nM Bafilomycin A1 and starved in EBSS for 2 hours prior to harvesting. **B)** Both WT and TD-ATG2 can recruit WIPI4 to LDs in HEK293 cells. Micrographs from fixed ATG2A/B double knockout cells transfected with WIPI-4-TurboGFP alone or in combination with 3xFlag-WT/TD-ATG2A-12xHis. ATG2A is visualized by immunofluorescence against its N-terminal 3xFlag tag, and LDs are visualized by BODIPY 558/568 C12 staining. **C)** TD-GFP-ATG2A binds to WIPI4. Immunoprecipitation (IP) of WT or TD GFP-ATG2A from ATG2 DKO cells stably expressing each construct using GFP Trap beads. WT HEK293 cells transiently transfected with EGFP served as a control for nonspecific binding. Each cell line was transiently transfected with WIPI-4-TurboGFP (not cross reactive with GFP Trap beads) **D)** Transport dead ATG2A does not drive a dominant negative autophagy phenotype in wildtype cells. LC3 flux and p62 clearance assays in indicated HEK293 cells performed identically to A. Ponceau stain (Total Protein) is shown as loading control.

## References

1. T. J. Melia, K. M. Reinisch, A possible role for VPS13-family proteins in bulk lipid transfer, membrane expansion and organelle biogenesis. J Cell Sci 135, (2022).

2. S. D. Neuman, T. P. Levine, A. Bashirullah, A novel superfamily of bridge-like lipid transfer proteins. Trends Cell Biol, (2022).

3. D. P. Valverde et al., ATG2 transports lipids to promote autophagosome biogenesis. J Cell Biol 218, 1787–1798 (2019).

4. N. Tamura et al., Differential requirement for ATG2A domains for localization to autophagic membranes and lipid droplets. FEBS letters 591, 3819–3830 (2017).

5. A. K. Velikkakath, T. Nishimura, E. Oita, N. Ishihara, N. Mizushima, Mammalian Atg2 proteins are essential for autophagosome formation and important for regulation of size and distribution of lipid droplets. Mol Biol Cell 23, 896–909 (2012).

6. S. Maeda, C. Otomo, T. Otomo, The autophagic membrane tether ATG2A transfers lipids between membranes. Elife 8, (2019).

7. T. Osawa et al., Atg2 mediates direct lipid transfer between membranes for autophagosome formation. Nat Struct Mol Biol 26, 281–288 (2019).

8. T. Osawa, Y. Ishii, N. N. Noda, Human ATG2B possesses a lipid transfer activity which is accelerated by negatively charged lipids and WIPI4. Genes Cells 25, 65–70 (2020).

9. Y. Hong, S. Li, J. Wang, Y. Li, In vitro inhibition of hepatic stellate cell activation by the autophagy-related lipid droplet protein ATG2A. Sci Rep 8, 9232 (2018).

10. E. Mailler et al., The autophagy protein ATG9A enables lipid mobilization from lipid droplets. Nat Commun 12, 6750 (2021).

11. H. Zhang et al., Proteome of skeletal muscle lipid droplet reveals association with mitochondria and apolipoprotein a-I. J Proteome Res 10, 4757–4768 (2011).

12. C. I. Pataki et al., Proteomic analysis of monolayer-integrated proteins on lipid droplets identifies amphipathic interfacial alpha-helical membrane anchors. Proc Natl Acad Sci U S A 115, E8172–E8180 (2018).

13. S. G. Pfisterer et al., Lipid droplet and early autophagosomal membrane targeting of Atg2A and Atg14L in human tumor cells. J Lipid Res 55, 1267–1278 (2014).

14. K. Bersuker et al., A Proximity Labeling Strategy Provides Insights into the Composition and Dynamics of Lipid Droplet Proteomes. Dev Cell 44, 97–112 e117 (2018).

15. A. Bieber et al., In situ structural analysis reveals membrane shape transitions during autophagosome formation. Proc Natl Acad Sci U S A 119, e2209823119 (2022).

16. S. Cai et al., In situ architecture of the lipid transport protein VPS13C at ER-lysosome membrane contacts. Proc Natl Acad Sci U S A 119, e2203769119 (2022).

17. P. Li, J. A. Lees, C. P. Lusk, K. M. Reinisch, Cryo-EM reconstruction of a VPS13 fragment reveals a long groove to channel lipids between membranes. J Cell Biol 219, (2020).

18. A. Bacle, R. Gautier, C. L. Jackson, P. F. J. Fuchs, S. Vanni, Interdigitation between Triglycerides and Lipids Modulates Surface Properties of Lipid Droplets. Biophys J 112, 1417–1430 (2017).

19. A. Chorlay, L. Foret, A. R. Thiam, Origin of gradients in lipid density and surface tension between connected lipid droplet and bilayer. Biophys J 120, 5491–5503 (2021).

20. Y. Wang et al., Construction of Nanodroplet/Adiposome and Artificial Lipid Droplets. ACS Nano 10, 3312–3322 (2016).

21. D. Hoekstra, N. Duzgunes, Lipid mixing assays to determine fusion in liposome systems. Methods Enzymol 220, 15–32 (1993).

22. N. Kumar et al., VPS13A and VPS13C are lipid transport proteins differentially localized at ER contact sites. J Cell Biol 217, 3625–3639 (2018).

23. A. Nguyen et al., Metamorphic proteins at the basis of human autophagy initiation and lipid transfer. Mol Cell 83, 2077–2090 e2012 (2023).

24. A. R. Thiam, R. V. Farese, Jr., T. C. Walther, The biophysics and cell biology of lipid droplets. Nat Rev Mol Cell Biol 14, 775–786 (2013).

25. A. Chorlay, A. Santinho, A. R. Thiam, Making Droplet-Embedded Vesicles to Model Cellular Lipid Droplets. STAR Protoc 1, 100116 (2020).

26. L. Caillon et al., Triacylglycerols sequester monotopic membrane proteins to lipid droplets. Nat Commun 11, 3944 (2020).

27. K. Ben M’barek et al., ER Membrane Phospholipids and Surface Tension Control Cellular Lipid Droplet Formation. Dev Cell 41, 591–604 e597 (2017).

28. R. Dabrowski, S. Tulli, M. Graef, Parallel phospholipid transfer by Vps13 and Atg2 determines autophagosome biogenesis dynamics. J Cell Biol 222, (2023).

29. R. Wu et al., High-resolution <em>de novo</em> structure prediction from primary sequence. bioRxiv, 2022.2007.2021.500999 (2022).

30. D. L. Brasaemle, N. E. Wolins, Isolation of Lipid Droplets from Cells by Density Gradient Centrifugation. Curr Protoc Cell Biol 72, 3 15 11–13 15 13 (2016).

31. S. Chowdhury et al., Insights into autophagosome biogenesis from structural and biochemical analyses of the ATG2A-WIPI4 complex. Proceedings of the National Academy of Sciences of the United States of America 115, E9792–E9801 (2018).

32. J. X. Zheng et al., Architecture of the ATG2B-WDR45 complex and an aromatic Y/HF motif crucial for complex formation. Autophagy 13, 1870–1883 (2017).

33. M. Bueno-Arribas et al., A conserved ATG2 binding site in WIPI4 and yeast Hsv2 is disrupted by mutations causing beta-propeller protein-associated neurodegeneration. Hum Mol Genet 31, 111–121 (2021).

34. T. J. Melia, A. H. Lystad, A. Simonsen, Autophagosome biogenesis: From membrane growth to closure. J Cell Biol 219, (2020).

35. J. Wang et al., An ESCRT-dependent step in fatty acid transfer from lipid droplets to mitochondria through VPS13D-TSG101 interactions. Nat Commun 12, 1252 (2021).

36. S. Chen et al., VPS13A and VPS13C Influence Lipid Droplet Abundance. Contact (Thousand Oaks) 5, 25152564221125613 (2022).

37. J. X. Tan, T. Finkel, A phosphoinositide signalling pathway mediates rapid lysosomal repair. Nature 609, 815–821 (2022).

